# VesiclePy: A Machine Learning Vesicle Analysis Toolbox for Volume Electron Microscopy

**DOI:** 10.1101/2025.09.08.674799

**Authors:** Jason K. Adhinarta, Yutian Fan, Adam Gohain, Michael Lin, Paige Nurkin, Richard Ren, Micaela Roth, Shulin Zhang, Ayal Yakobe, Rafael Yuste, Donglai Wei

**Author notes:** These authors contributed equally to this work (Jason K. Adhinarta, Yutian Fan, Adam Gohain, Michael Lin, Paige Nurkin, Richard Ren, Micaela Roth, Shulin Zhang).

## Abstract

Vesicles are critical components of neurons that package neurotransmitters and neuropeptides for their release, in order to communicate with other neurons and cells. However, due to their small size, the reconstruction of the full vesicle endowment across an entire neuronal morphology remains challenging. To achieve this, we have used, as a tool to identify and visualize vesicles, Volume Electron Microscopy (vEM), a method that has the nanoscale resolution to detect individual vesicle boundaries, content, and 3D locations. However, the large volume of vEM datasets poses a challenge in the segmentation, classification, and spatial analysis of tens of thousands of vesicles and their target cell in 3D. Here we report the development of VesiclePy, an integrated pipeline for automated segmentation, classification, proofreading, and spatial analysis of vesicles, relative to neuron masks in large-volume electron microscopy data. Our package integrates the efficiency of deep learning and the accuracy of human proofreading and provides a streamlined package in chunked processing and accurate indexing, localization, and visualization of single vesicle resolution in large vEM data. We demonstrate the viability of VesiclePy using high-pressure frozen serial EM data of *Hydra vulgaris* and quantify the performance of the package using ground truth manual annotations. We show that VesiclePy can process a multiterabyte serial EM dataset, efficiently annotate 53,851 vesicles from 20 complete neurons, and classify vesicles into 5 types. Each vesicle has a unique ID and 3D location for further spatial analysis in relation to neuron or non-neuronal targets nearby. Finally, by combining vesicle data and morphological information of each neuron, we can quantitatively cluster neurons into subtypes. VesiclePy is available at https://github.com/PytorchConnectomics/VesiclePy under an MIT license.

## Introduction

Neurons use vesicles to store neurotransmitters and neuropeptides and release them at the membrane to communicate with neighboring and distant cells. Understanding vesicle morphology and spatial distribution helps understand the content, mechanism, and target of neuronal communication. Given their small size and transient nature, electron microscopy is well-suited for capturing sub-cellular resolution vesicle membranes and structures. However, most analyses are limited to synapses and small vesicles using manual [1, 2], or automated [3, 4] segmentation, and large vesicles containing neuropeptides and their release site outside synapses have not been systematically characterized in 3D in full neurons. Here we present VesiclePy, a Python pipeline that provides a streamlined process for automated segmentation, classification, proofreading, spatial analysis, and visualization of vesicles, as well as the classification of neurons according to vesicle composition and neuron morphology. In this example dataset, we obtained from a *Hydra* 1,829 sections of serial electron microscopy (SEM) images and 20 neuron segmentations in half of the endoderm [5]. *Hydra* has a simple nervous system of a few hundred neurons, and a sparsely distributed nerve net with few neuronal contacts (Fig 1A). *Hydra*’s endodermal neurons contain a great number of large vesicles, including dense core vesicles (DCV), clear vesicles (CV, Fig 1B), dense core vesicles with halo (DCVH, Fig 1C), each of which may contain neuropeptides and neurotransmitters. There are also small clear vesicles (SCV) and small dense vesicles (SDV), mostly found in soma, and may not participate in synaptic release. The location of vesicles can inform us about their potential target, in this case, usually not near another neuron, suggesting asynaptic paracrine release. Each neuron contains all types of vesicles, however, the proportion of each vesicle type may differ across neuron types.

**Fig 1.**
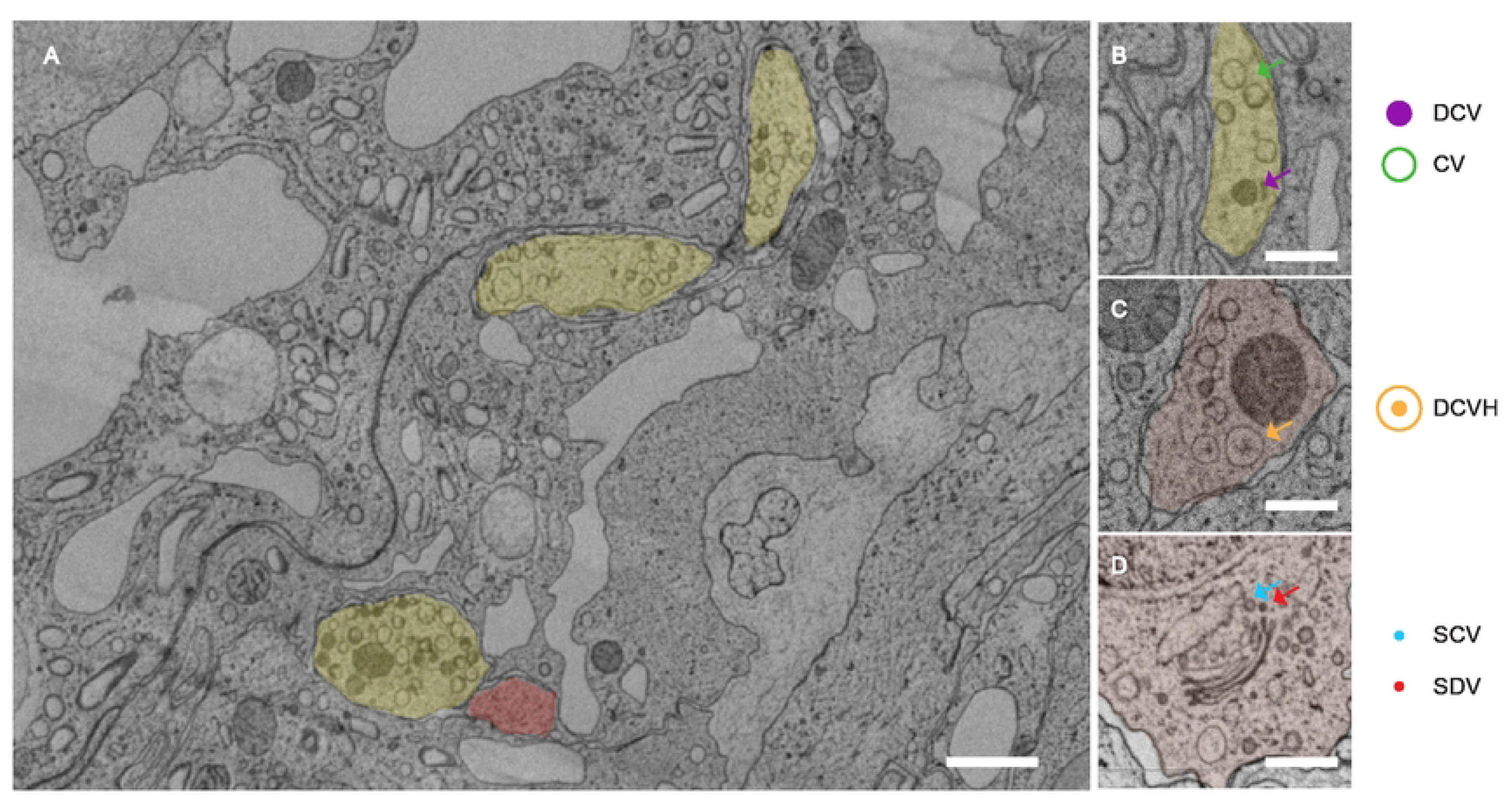
EM image, neuron segmentation and five types of vesicles in *Hydra Vulgaris*. A) Representative EM section, with yellow segmentation over one neuron and red over another neuron, Scale bar 1 μm (B) A closeup EM image of dense core vesicles (DCV, purple arrow) and clear vesicles (CV, green arrow) in a neuron (yellow). (C) Two dense core vesicles with halo (DCVH, orange arrow) in a neuron (red). (D) Small clear vesicles (SCV, blue arrow) and small dense vesicles (SDV, red arrow) in the soma near the Golgi apparatus. (B)-(D) Scale bar 500 nm.

### Design and Implementation

We designed the pipeline that processes large and small vesicles in parallel, with modular processing centres for segmentation, classification, analysis, visualization, and additional statistical and clustering analysis. Segmentation consists of an iterative deep learning model to automatically segment large vesicles from ground truth data, and manual segmentation of small vesicles. Classification of vesicle types consists of a supervised deep learning model based on ground truth on large vesicle types, and an unsupervised autoencoder and classification of small vesicle types. Analysis produces morphology information and 3D location of each vesicle, this information can further be used to compute the vesicle density and vesicle intersection with regions of interest.

Finally, we can produce visualization on multiple resolution scales for different purposes, as well as statistics and clustering of neurons based on vesicle and neuron morphology information (Fig 2).

**Fig 2.**
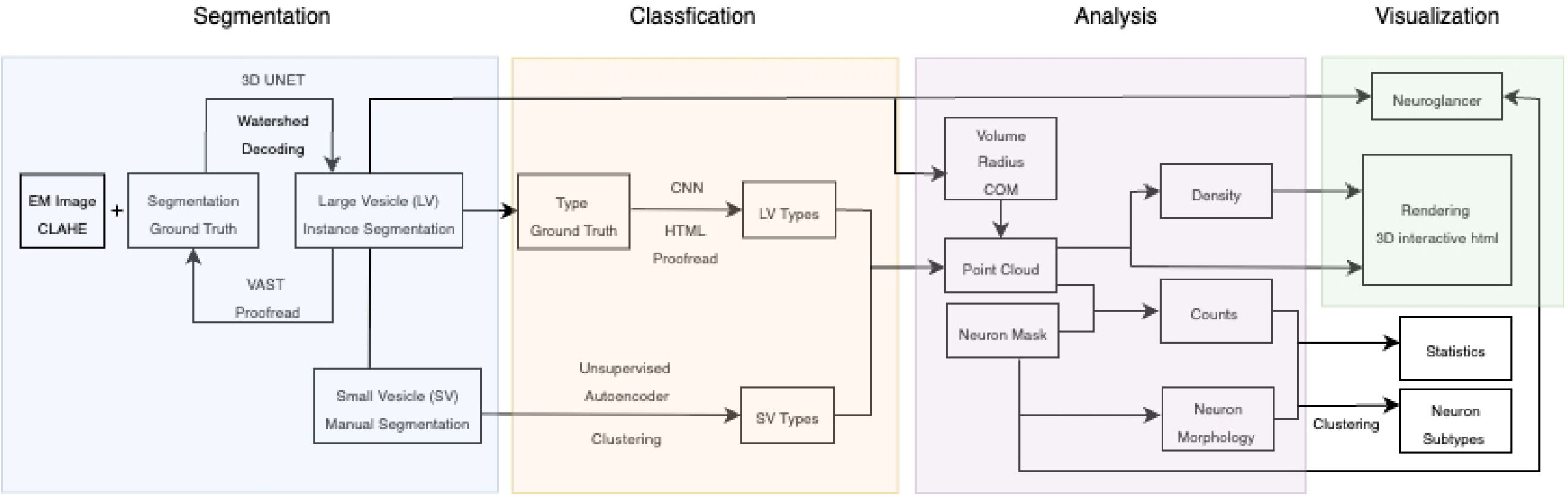
Pipeline overview. Flowchart of the computational pipeline in 5 modules.

### Large Vesicle Instance Segmentation

Vesicle detection and segmentation of vesicles individually in 3D is the foundation for any downstream analysis on vesicle morphology and locations. However, every neuron has thousands of sparsely distributed vesicles, most of which span multiple z-slices.

Human annotation would be extremely labor-intensive and prone to errors. Thus, we used an iterative semi-automated 3D-UNet [6] convolutional neural network, and specifically the architecture described in the NucMM challenge (19), that learns from user-annotated volumes and provides multichannel 3D semantic segmentation predictions. Furthermore, we used CLAHE [7] as a preprocessing step to increase image contrast, and then applied multichannel watershed to achieve instance segmentation of individual vesicles. These predictions are further corrected by humans and learned by the model to produce better predictions. Additionally, digital enhancement of the image and sampling of erroneous regions further improved the prediction result. Overall, this pipeline was able to efficiently produce highly accurate instance segmentation of every single large vesicle in all neurons in 3D.

### Iterative training and prediction

Development followed a cyclical process of training, prediction, human correction, and sampling of erroneous regions to improve the model’s performance. Firstly, we annotated multiple ground truth volumes and separated them into training or testing partitions. Then, the model was trained on the training partition and generated predictions on the testing partition. Upon proofreading and correction of the prediction result, we identified failure patterns and selected additional areas that targeted patterns that needed improvement. We then generated predictions on these volumes and corrected any errors so that they could be used as ground truth in the next cycle of training and prediction. For example, after two iterations of the training cycle using 26 volumes, 340,092,727 voxels of training data (Fig 3D-E), we found poor qualitative performance in regions containing a high concentration of dense core vesicles and dense core vesicles with a halo. Using connected components, we find 33 different small subvolumes, 354,022,199 voxels that have a concentration of these dense vesicles, then manually correct them. These subvolumes are used as training data in the third training interaction (Fig 3F).

**Fig 3.**
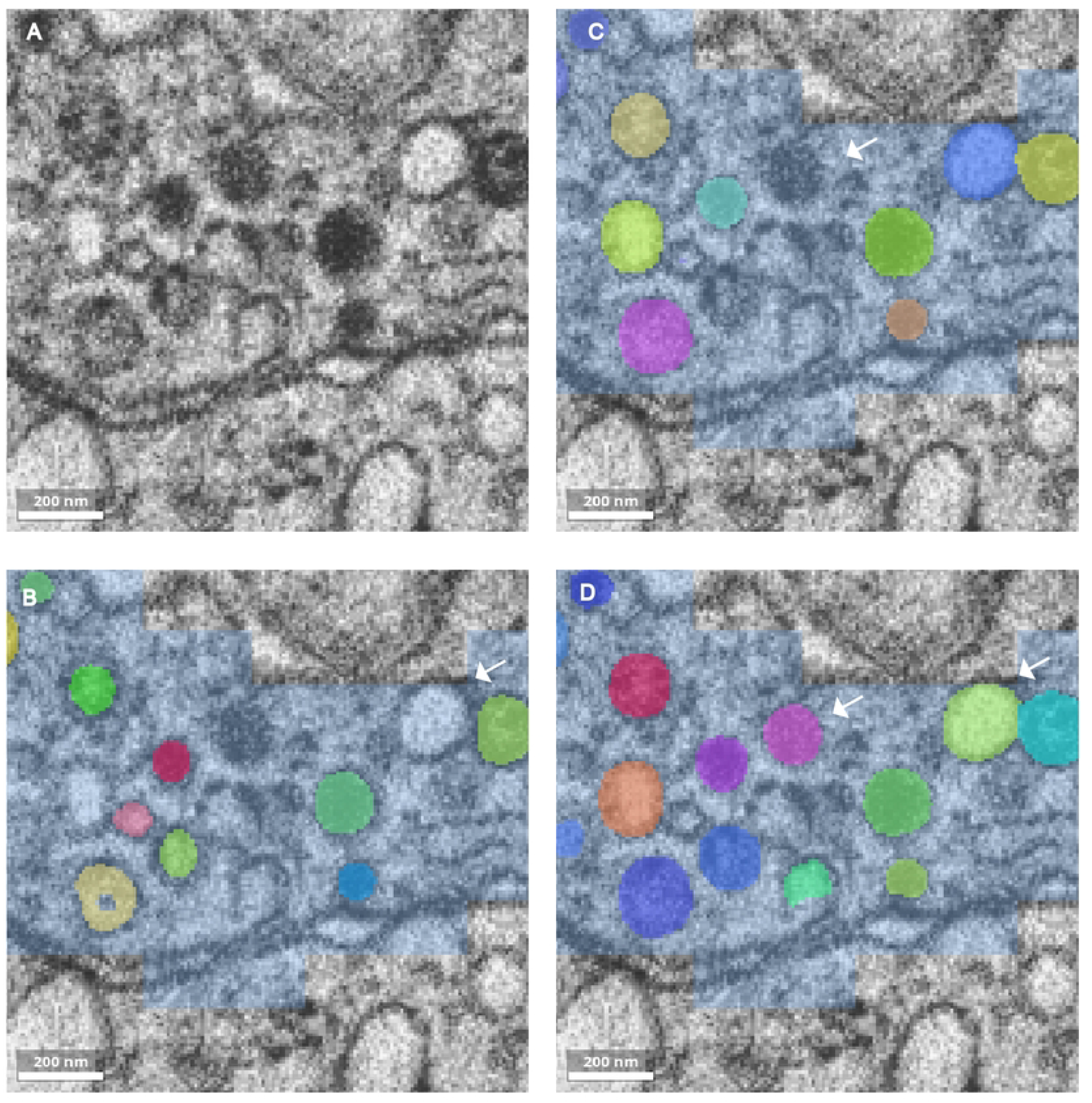
Example result at each training checkpoints. (A) EM image after applying CLAHE (B) Example of prediction result on CLAHE processed image, after 100,000 epochs of training, white arrow marks a false negative vesicle (C) Result after 300,000 epochs of training, white arrow marks a false negative dense core vesicle (D) Result after 1,000,000 epochs of training, white arrows mark improved result that was previously false negative. Scale bar (A)-(D) 200 nm.

### Instance segmentation

In order to achieve instance segmentation of vesicles, each with a unique segmentation ID. We chose to predict a three-channel volume consisting of binary segmentation, contour, and distance transform predictions. We then used contour and distance transform predictions to generate seeds for watershed segmentation [8], which detected the boundary of each vesicle. Finally, we collapsed the three-channel prediction into a single-channel prediction of individual vesicles, each with unique segmentation IDs.

### Supervised Vesicle Type Classification

Morphology of vesicles is useful in determining their type, content, and state in a dynamic biological process. In the example dataset, we could identify three morphologically distinct large vesicle types as previously described: CV(Fig 4A), DCV (Fig 4B), and DCVH (Fig 4C). Each vesicle spans multiple z-sections, sometimes with varied appearances on different sections. We used a 3D convolutional neural network [9] to automate the classification process. We also developed an HTML-based proofreading tool that displayed the results of vesicles of the same kind on one page, which allowed for efficient human proofreading and correction. The network processes 31×31×5 voxel stacks using a series of convolutional layers with batch normalization and ReLU activations, followed by max pooling and fully connected layers to produce categorical predictions. Despite its architectural simplicity, the network captured essential morphological features relevant to vesicle classification. To mitigate class imbalance and improve generalization, we constructed a balanced training dataset by aggregating vesicles from all available manually proofread cells and sampling 1916 of each CV, DCV, and DCVH examples. When trained on this dataset and evaluated on a held-out collection of previously labeled data, the model achieved a validation accuracy of 83.6%. The evaluation result (Table 2) showed that the network was able to distinguish between vesicles of very distinct features (DCV and CV). However, vesicles with mixed morphology (DCVH) had a lower performance.

**Table 1.** Segmentation model training and performance.

| Training Epochs | RAND | Precision | Recall | Example Result |
| --- | --- | --- | --- | --- |
| 100,000 | 0.3446 | 0.9914 | 0.4982 | Fig 3A |
| 300,000 | 0.16242 | 0.9848 | 0.7340 | Fig 3B |
| 1,000,000 | 0.1517 | 0.9842 | 0.7492 | Fig 3C |
| 1,000,000 | 0.2220 | 0.9798 | 0.6516 | Fig 3D |

Evaluation of model performance in RAND, precision and recall scores, and corresponding example result at each checkpoint.

**Table 2.** Classification model performance on different vesicle types.

| Type | RAND | Precision | Recall |
| --- | --- | --- | --- |
| CV | 0.3125 | 0.6931 | 0.6820 |
| DCV | 0.0991 | 0.9099 | 0.8921 |
| DCVH | 0.8655 | 0.0941 | 0.2353 |
| Total | 0.4162 | 0.5657 | 0.6031 |

Evaluation of the model performance of each type of large vesicle and in total in RAND, precision and recall scores.

**Fig 4.**
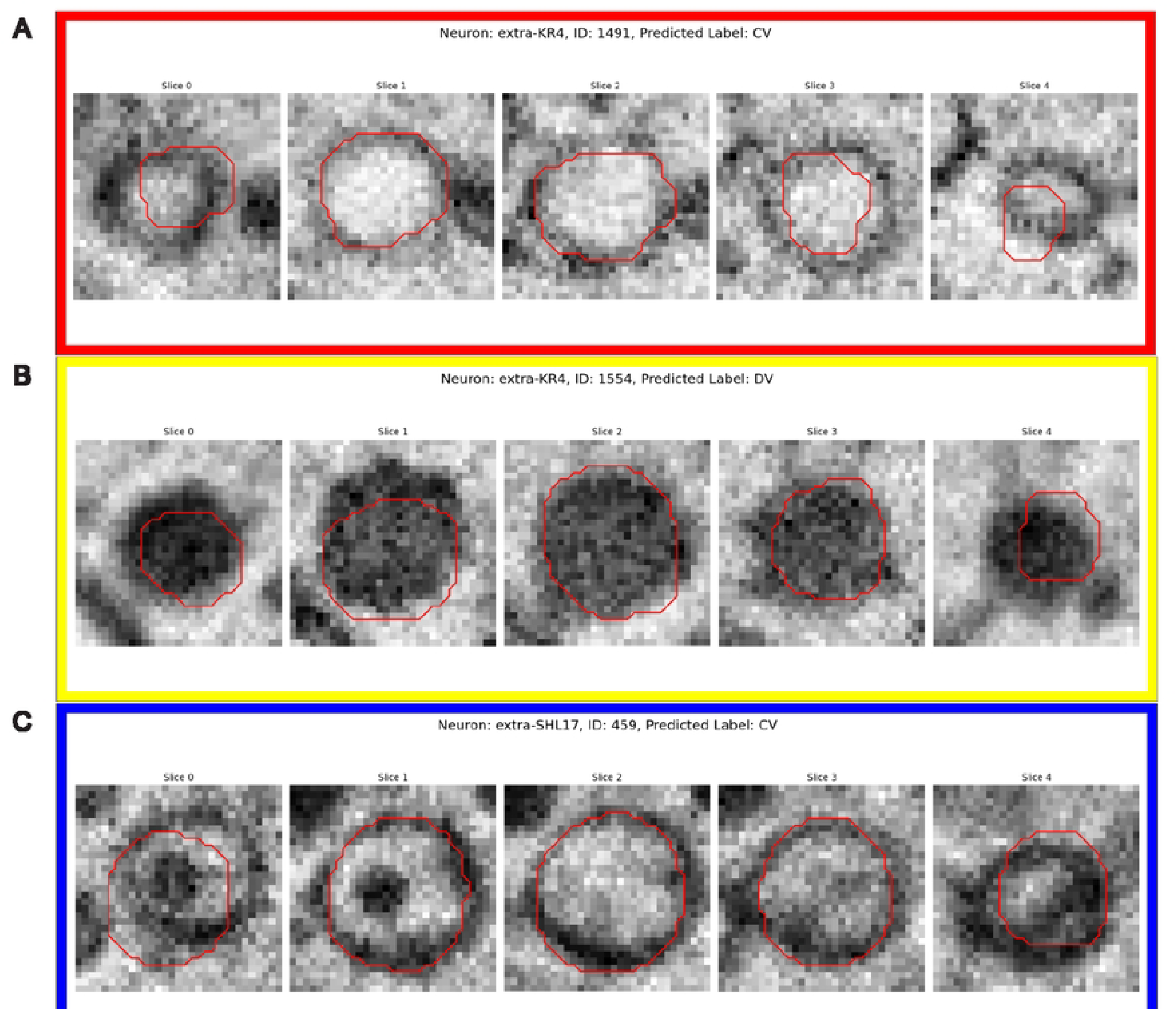
Vesicle type classification proofreading interface. Examples of HTML-based proofreading pages from each vesicle type. (A) 5 Z-stack images of CV with red outlines of the vesicle boundary, bounded by a red box. (B) 5 Z-stacks of DCV, bounded by a yellow box. (C) 5 Z-stacks of DCVH, bounded by a blue box.

To ensure the robustness of the pipeline, we implemented an iterative workflow that integrates classification predictions with visualization and manual proofreading in HTML format. Input data consisted of 3D vesicle image stacks and their corresponding bounding box files, organized into neuron-specific subdirectories. The pipeline first applies the trained network to classify each vesicle and outputs its prediction along with a visualization image. These results are organized into category-specific subfolders (CV, DCV, DCVH), providing users with an overview of the classification result that can be proofread and corrected by simply clicking.

### Unsupervised Small Vesicle Type Classification

Small vesicles are operationally defined as vesicles with a diameter of less than 80 nm; they are much smaller than the large vesicles and difficult to classify manually by morphology. Instead of training a supervised classification model to predict vesicle types, as we did in the previous section, here we performed unsupervised dimensionality reduction to determine whether distributional analysis (made feasible on a lower number of dimensions) can be used to naturally reveal clusters of vesicle types.

To ensure accurate vesicle identification, we manually annotated segmentations for the small vesicles (Fig 5A). We extracted image patches centered around vesicle bounding boxes and resized the patches to 11×11 pixels to standardize differing vesicle shapes and sizes. To account for potential translations and rotations in vesicle appearances, we utilized variational autoencoders with in-built translation and rotation invariance, provided by the pyroVED [10] library, allowing the model to be robust to small segmentation errors, which may cause the vesicles to be off-center. Using this model, each 11×11 image patch is compressed into an embedding with two latent dimensions. We plotted predicted vesicle appearances according to the corresponding latent coordinates (Fig 5B), and a heatmap of the embeddings (Fig 5C). Finally, to cluster the density map into two categories (determined by visual inspection) we used the HDBSCAN [11] algorithm as it is a standard approach for clustering low-dimensional measurements with noise (Fig 5D).

**Fig 5.**
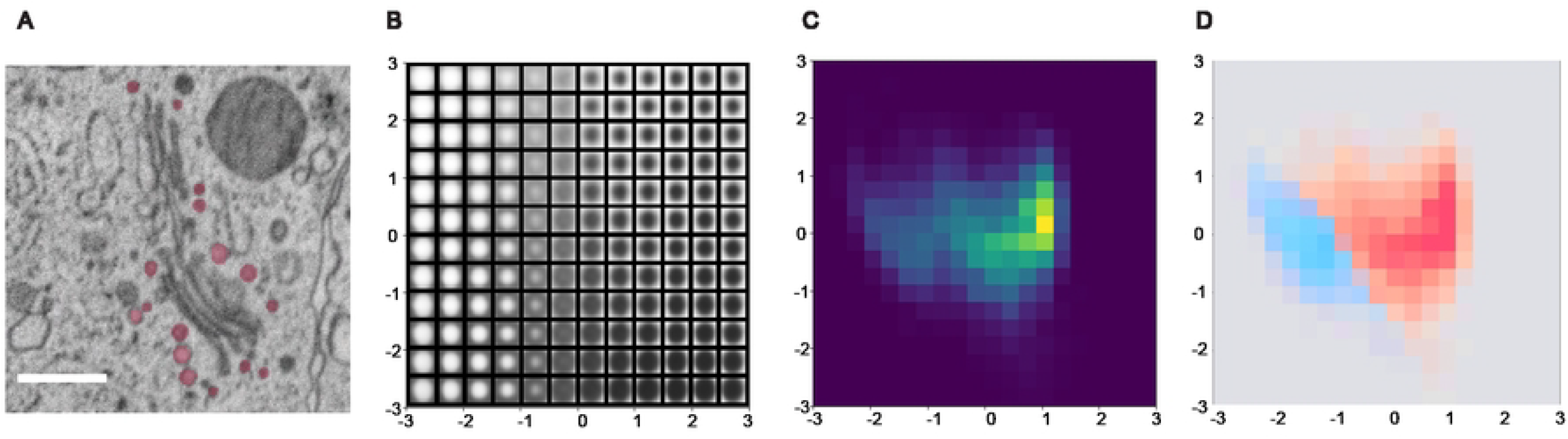
Unsupervised small vesicle type classification. (A) Manual semantic segmentation of small vesicles. Scale bar 500 nm (B) Variational autoencoder classifications (C) Embedding distribution heatmap (D) HDBSCAN clustering of vesicle types with density map.

### Morphology and Spatial Analysis

Unlike small vesicles, large and dense core vesicles are often released in a paracrine manner outside of synapses ([5, 12]). Analyzing their spatial distribution and potential targets requires examining large regions spanning the entire neuron and its neighbors. To manage this computationally intensive task, we confined analysis to minimal bounding boxes per neuron, stored in high-resolution HDF5 format. Vesicle data within these regions were converted into point clouds that retain key morphology, spatial, and segmentation metadata for efficient downstream processing.

### Neuron mask usage

To identify regions of possible vesicle release that target another neuron, we stitched adjacent neurons into unified bounding boxes using global coordinates, and identified regions “near” another neuron by computing a Euclidean Distance Transform [13] from the stitched pieces—based on a threshold determined by the average vesicle diameter plus 2 times standard deviation (Fig 6). To account for segmentation uncertainty, we added 1–3 voxel buffers around both source and adjacent neurons, then extracted surface patches within these margins. The voxel counts of these regions were scaled by resolution to compute surface area.

**Fig 6.**
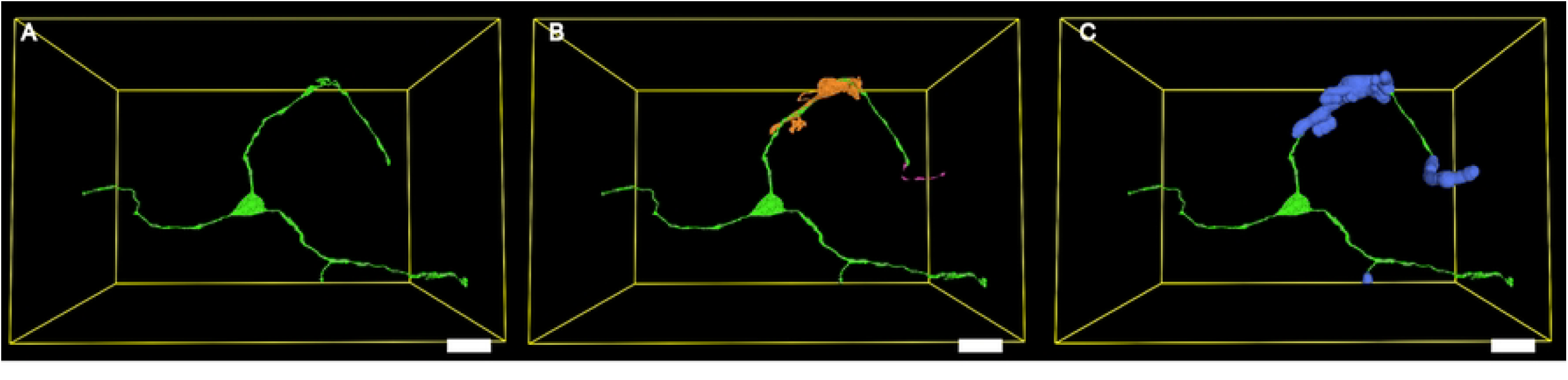
Neighbouring neuron and potential vesicle release region. (A) 3D rendering of one neuron and its minimal bounding box. (B) Neuron and adjacent neuron pieces that are within the bounding box. (C) Expanded the adjacent neuron mask that determines the “near another neuron” region in the target neuron. (A) - (C) Scale bar 10 μm.

### Vesicle morphology

The high resolution needed to compute vesicle morphology and the large number of vesicles in each neuron demand extensive computational resources. Therefore, we divided the task into three parts, each leveraging the appropriate file format. Firstly, we computed the morphology of each vesicle using the original segmentation masks. Vesicle volumes were calculated by summing segmented voxels and scaling by resolution. Radii were estimated by collapsing each roughly spherical mask along its lowest-resolution axis and measuring the equivalent circular area. Secondly, vesicles were converted into point clouds storing their center of mass (COM), volume, radius, classification type, and ID. This format allowed for compact exports and efficient analysis using tools like Pandas [14] and Polars [15], which are poised to process column-based data. Finally, vesicle distribution densities were calculated instance-wise using kd-trees [16], undergoing a nearest neighbors query within 500 nm of each vesicle COM. These values were normalized across the dataset to visualize relative density gradients.

### Counting vesicles

Finally, to determine the number of vesicles within a specific neuronal region of interest, we extracted values of a binary mask of the region from all of the stored COM coordinates of the vesicles within the neuron, thus avoiding direct use of the original high-resolution vesicle segmentation data. For example, we calculated the number of vesicles present in “near-neuron” regions by using distance thresholds to extract masks of these regions, then utilizing the point cloud metadata to keep track of vesicle type distributions. Similarly, we calculated the number of small vesicles present within soma regions to determine whether there was a greater density of small vesicles within the soma.

### Visualization

Effective visualization remains a persistent challenge in the analysis of large-scale volume electron microscopy (vEM) datasets [17], where the complexity and density of biological structures require tailored tools for rendering, inspection, and semantic interpretation. To support efficient human-in-the-loop data analysis and observational hypothesis generation, we implement a modular visualization pipeline built on four complementary platforms—Neuroglancer [18], PyVista [19], Plotly [20], and three.js [21]—each addressing specific bottlenecks in interoperability, scalability, and semantic clarity.

Neuroglancer (Fig 7A) anchors the workflow by allowing direct inspection of EM data and segmentation overlays for ground truth validation. Although its format limitations restrict customization (e.g., dynamic color encodings or metadata overlays), it anchors all downstream visualization through persistent vesicle ID links across tools. PyVista (Fig 7B) enables more flexible, high-fidelity mesh visualization, where features like vesicle subtype or spatial density can be encoded directly as colormaps. However, PyVista’s performance degrades on full-volume datasets and requires substantial preprocessing, limiting its scalability for larger data exploration tasks. Plotly (Fig 7C), allows for browser-native 3D rendering of vesicle point clouds and mesh overlays, with hover-based metadata inspection and standalone HTML export. This allows for quick inspection of individual neurons or vesicle subsets, but struggles to scale to full datasets due to computational constraints. For dataset-wide visualization at scale, we develop a lightweight three.js-based viewer (Fig 6D) optimized for rendering tens of thousands of vesicles and neurons with subtype and density encodings. Though it lacks raw image overlays, it fully supports metadata and links back to Neuroglancer, preserving semantic disambiguation without sacrificing rendering performance.

**Fig 7.**
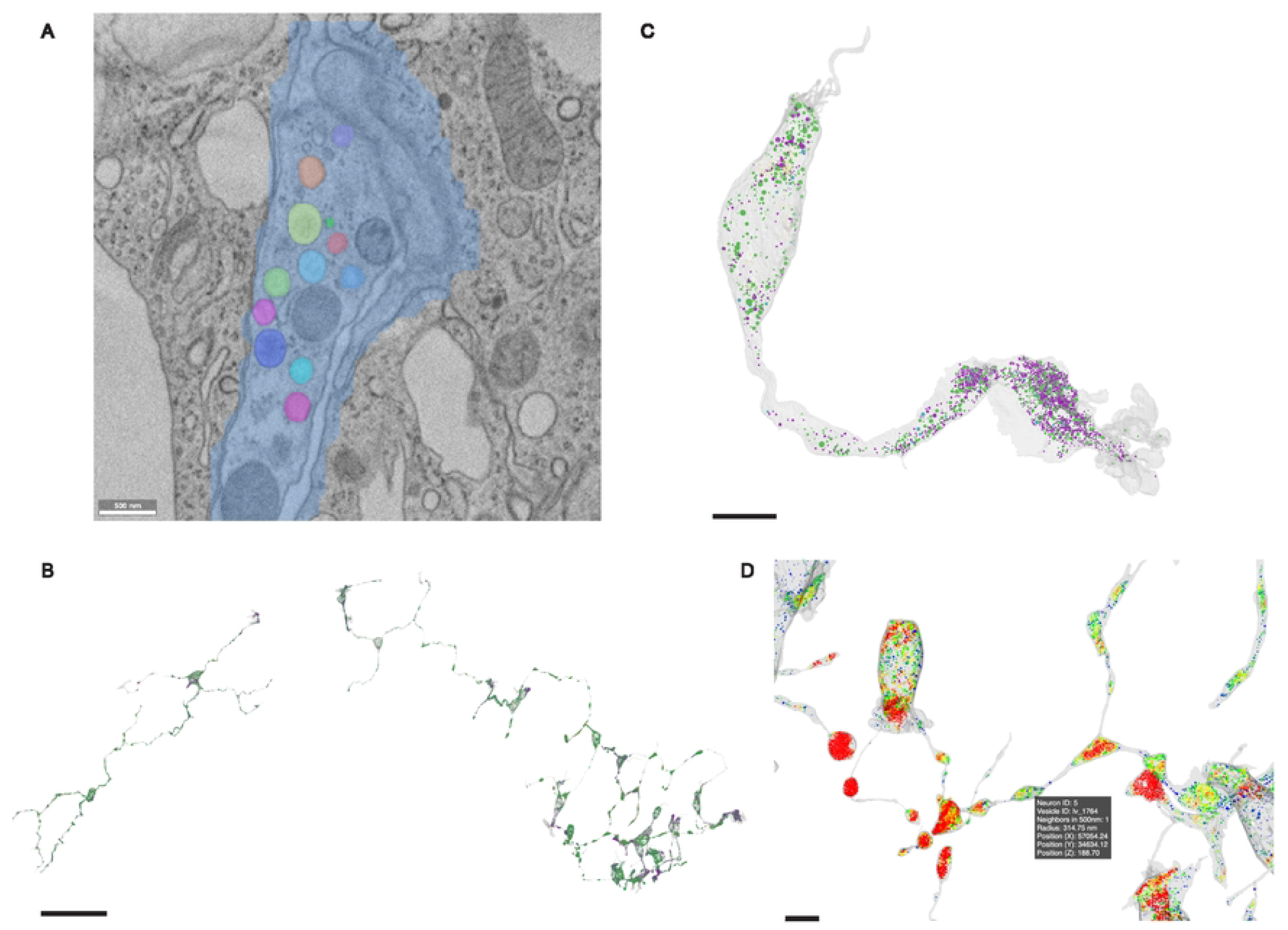
A four-platform visualization pipeline for vEM data analysis. A four-platform visualization pipeline for vEM data analysis. (A) Neuroglancer anchors ground truth validation via EM overlays and segmentation. Scale bar 500 nm. (B) PyVista supports high-fidelity local mesh rendering with colormaps, e.g., for subtype and density. Scale bar 50 μm. (C) Plotly enables interactive browser-based mesh and point cloud inspection. Scale bar 5 μm. (D) Three.js provides scalable full-dataset visualization with metadata support. Scale bar 5 μm.

### Neuron Type Cluster Analysis

Neuron morphology and connectivity are common parameters for clustering of neuron types in connectomics studies [22, 23]; however, when connectivity extends beyond physical synaptic connections, we propose the combination of morphology and vesicles as additional valid clustering parameters. In this example dataset, an initial categorization of 20 neurons into 5 distinct clusters was established from distinctive morphology features [5]. To further validate these clusters, we utilized both morphology and vesicle information in an unsupervised classification method. Vesicle compositions of each neuron from the previous analysis, and several categories of morphology data (polarity, cilia, microvilli, handshake, volume, and length) for the neurons were extracted. Numerical features were logarithmically normalized from [0,1]. Subsequently, the Gower distance metric was calculated from this processed mixed-data feature set, producing a 20×20 pairwise dissimilarity matrix. Gower distance provides a unified similarity measure across these diverse feature types, enabling a holistic comparison relevant to overall neuron structure [24]. Using this Gower matrix, hierarchical clustering with the ‘complete’ linkage method was performed [25]. Cutting the resulting dendrogram at a distance threshold of 0.4 yielded 4 distinct neuron clusters (Fig 8) that corresponded with 4 clusters established through qualitative morphology classification, and 1 cluster (Neuron 10) merged.

**Fig 8.**
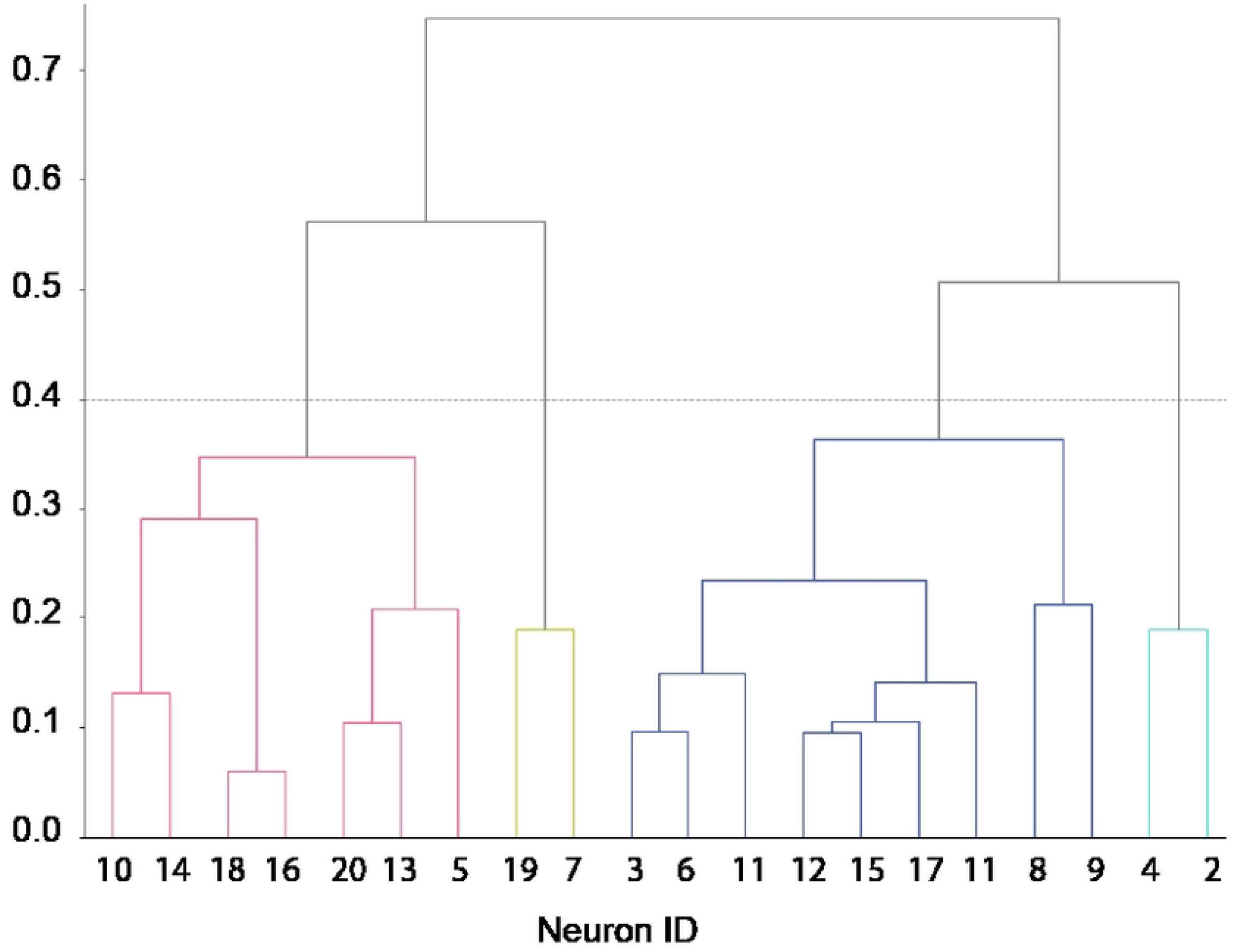
Neuron Type Cluster Analysis. A dendrogram of neuron clustering based on Gower’s distance, with a distance threshold at 0.4.

## Results

The example dataset is an SEM taken from a high-pressure frozen *Hydra vulgaris*. The pipeline segmented a total of 53,851 vesicle instances, including 42,235 large vesicles and 11,616 small vesicles in 20 complete segmented neurons. We classified all large vesicles into these three categories, yielding 24,998 CV, 15,188 DCV, and 2049 DCVH. For Small vesicles, we classified 2305 SCV and 9311 SDV. Next, we extracted the volume and diameter of each vesicle and computed the overlap between the vesicle coordinates with the “near” and “far” from another neuron region, and quantified the percentage of vesicles “near” another neuron is significantly less than “far” across different neuronal types and vesicle types. We also created 2D rendering and 3D interactive HTML for visualization and inspection of all vesicle types, their location, and density in relation to the neuron mask. Finally, using vesicle count by type for each neuron and additional morphological measurements and features, we were able to cluster neurons into 4 subtypes that aligned with our initial qualitative classification based on morphology.

## Acknowledgments

The project is supported by the NSF (CRCNS 1822550; 2203119; 2239688) and the Vannevar Bush Faculty Award (ONR N000142012828).

## Author Contribution

S.Z, D.W. and R.Y. designed and conceptualized the project. J.A., Y.F., A.G., M.L., R.R., M.R., A.Y. designed and developed the software pipeline. P.N. proofread results. S.Z., J.A., Y.F., A.G., M.L., R.R., M.R., D.W. and R.Y wrote the initial manuscript and edited the paper. D.W. and R.Y directed the project and secured funding.

## References

1. Khanmohammadi M, Waagepetersen R, Sporring J. Analysis of shape and spatial interaction of synaptic vesicles using data from focused ion beam scanning electron microscopy (FIB-SEM). Frontiers in Neuroanatomy. 2015;9. doi:10.3389/fnana.2015.00116.

2. Schindelin J, Arganda-Carreras I, Frise E, Kaynig V, Longair M, Pietzsch T, et al. Fiji: an open-source platform for biological-image analysis. Nature Methods. 2012;9(7):676–682. doi:10.1038/nmeth.2019.

3. Khosrozadeh A, Seeger R, Witz G, Radecke J, Sørensen JB, Zuber B. CryoVesNet: A dedicated framework for synaptic vesicle segmentation in cryo-electron tomograms. Journal of Cell Biology. 2024;224(1):e202402169. doi:10.1083/jcb.202402169.

4. Muth S, Moschref F, Freckmann L, Mutschall S, Hojas-Garcia-Plaza I, Bahr JN, et al. SynapseNet: Deep Learning for Automatic Synapse Reconstruction; 2024. Available from: http://biorxiv.org/lookup/doi/10.1101/2024.12.02.626387.

5. Zhang S, Ofer N, Dupre C, Schalek R, Wu Y, Lin M, et al. Ultrastructural reconstruction of the endodermal nerve net of Hydra vulgaris; 2025.

6. Özgün Cicek, Abdulkadir A, Lienkamp S, Thomas B, Ronneberger O. 3D U-Net: Learning Dense Volumetric Segmentation from Sparse Annotation; 2016. Available from: https://arxiv.org/abs/1606.06650.

7. Zuiderveld K. Contrast limited adaptive histogram equalization. In: Graphics gems IV. USA: Academic Press Professional, Inc.; 1994. p. 474–485.

8. He J, Zhang S, Yang M, Shan Y, Huang T. Bi-Directional Cascade Network for Perceptual Edge Detection; 2019. Available from: http://arxiv.org/abs/1902.10903.

9. Yang J, Shi R, Wei D, Liu Z, Zhao L, Ke B, et al. MedMNIST v2 - A large-scale lightweight benchmark for 2D and 3D biomedical image classification. Scientific Data. 2023;10(1):41. doi:10.1038/s41597-022-01721-8.

10. Ziatdinov M. pyroVED: Variational encoder-decoder for unsupervised learning in Pyro; 2020. Available from: https://github.com/ziatdinovmax/pyroVED.

11. McInnes L, Healy J, Astels S. hdbscan: Hierarchical density based clustering. Journal of Open Source Software. 2017;2(11):205. doi:10.21105/joss.00205.

12. Zhu PC, Thureson-Klein A, Klein RL. Exocytosis from large dense cored vesicles outside the active synaptic zones of terminals within the trigeminal subnucleus caudalis: A possible mechanism for neuropeptide release. Neuroscience. 1986;19(1):43–54. doi:10.1016/0306-4522(86)90004-7.

13. Silversmith W. edt: Euclidean Distance Transform for Python; 2023. Available from: https://pypi.org/project/edt/.

14. team Tpd. pandas-dev/pandas: Pandas; 2020. Available from: 10.5281/zenodo.3509134.

15. Vink R, de Gooijer S, Beedie A nameexhaustion, Gorelli ME, Burghoorn G, et al. pola-rs/polars: Rust Polars 0.49.1; 2025. Available from: https://zenodo.org/doi/10.5281/zenodo.7697217.

16. Virtanen P, Gommers R, Oliphant TE, Haberland M, Reddy T, Cournapeau D, et al. SciPy 1.0: fundamental algorithms for scientific computing in Python. Nature Methods. 2020;17(3):261–272. doi:10.1038/s41592-019-0686-2.

17. Müller A, Schmidt D, Albrecht JP, Rieckert L, Otto M, Galicia Garcia LE, et al. Modular segmentation, spatial analysis and visualization of volume electron microscopy datasets. Nature Protocols. 2024;19(5):1436–1466. doi:10.1038/s41596-024-00957-5.

18. Maitin-Shepard J, others. Neuroglancer: Web-Based Visualization of Large-Scale 3D Electron Microscopy Data. In: Conference on Neural Information Processing Systems (NeurIPS); 2017. Available from: https://github.com/google/neuroglancer.

19. Sullivan C, Kaszynski A. PyVista: 3D plotting and mesh analysis through a streamlined interface for the Visualization Toolkit (VTK). Journal of Open Source Software. 2019;4(37):1450. doi:10.21105/joss.01450.

20. Inc PT. Plotly: Collaborative data science; 2015. Available from: https://plotly.com/python/graph-objects/.

21. Cabello R. Three.js: JavaScript 3D Library; 2010. Available from: https://threejs.org.

22. Schlegel P, Yin Y, Bates AS, Dorkenwald S, Eichler K, Brooks P, et al. Whole-brain annotation and multi-connectome cell typing of Drosophila. Nature. 2024;634(8032):139–152. doi:10.1038/s41586-024-07686-5.

23. Liu L, Yun Z, Manubens-Gil L, Chen H, Xiong F, Dong H, et al. Connectivity of single neurons classifies cell subtypes in mouse brains. Nature Methods. 2025;22(4):861–873. doi:10.1038/s41592-025-02621-6.

24. On the challenge of treating various types of variables: application for improving the measurement of functional diversity - Pavoine - 2009 - Oikos - Wiley Online Library;. Available from: https://nsojournals.onlinelibrary.wiley.com/doi/abs/10.1111/j.1600-0706.2008.16668.x.

25. Jay JJ, Eblen JD, Zhang Y, Benson M, Perkins AD, Saxton AM, et al. A systematic comparison of genome-scale clustering algorithms. BMC bioinformatics. 2012;13 Suppl 10(Suppl 10):S7. doi:10.1186/1471-2105-13-S10-S7.

